# A meta-analysis on the benefits and costs of hosting secondary endosymbionts in sap-sucking insects

**DOI:** 10.1101/563031

**Authors:** Sharon E. Zytynska, Karim Thighiouart, Enric Frago

## Abstract

Herbivorous insects host various bacteria that help them to feed, grow, and survive. Sap-sucking insects, in particular, feed on a nutrient-poor resource and have evolved obligate symbioses with nutritional bacteria for survival. Additionally, sap-sucking insects have formed facultative associations with bacterial symbionts that are not essential for growth and survival but assumed to confer some benefits, such as resistance to natural enemies. Several recent reviews have highlighted the importance of these symbionts in understanding their hosts’ biology, but currently there is a lack of a quantitative and systematic analysis of the published evidences exploring whether the different endosymbionts are actually beneficial or not. In this meta-analysis we explored the potential costs and benefits associated with hosting facultative endosymbionts in sap-sucking insects. Our first result is that most of the empirical experimental data information is limited to a few species of aphid and one species of whiteflies. Through the meta-analysis we showed that hosting symbionts generally leads to costs through increased development time, reduced longevity, and reduced fecundity, and benefits via increased resistance to parasitic wasps in sap-sucking insects. However, the impact of these costs and benefits was strongly insect and symbiont species dependent. Many of the insects studied are agricultural pests, and understanding the impact of bacterial symbionts on their hosts across different environments can benefit sustainable management of greenhouses and agricultural land.

## Introduction

Insect associations with mutualistic microbes are widespread, and it is nowadays widely accepted that these symbionts, and bacteria in particular, play key roles in the biology of their hosts (Brownlie & Johnson 2009; Feldhaar 2011). Thanks to important innovations in molecular techniques, the last two decades have provided deep insights into these diverse and often intricate interactions. The guts of most insects that feed on plant leaves, for instance, are colonised by complex communities of bacteria and fungi (Dillon & Dillon 2004). The role played by most of these microbes has yet to be understood, but evidence suggests that they are an important component of the hosts immune system, and that they often assist in the digestive process. Relative to leaf-feeding insects, the nutritional services provided by bacterial symbionts in sap-sucking insects are better understood (Moran, McCutcheon & Nakabachi 2008). This lifestyle has evolved multiple times among Hemiptera and includes most Sternorrhyncha (whiteflies, mealybugs, aphids and psyllids), many Auchenorrhyncha (planthoppers and leafhoppers) and most herbivorous Heteroptera (lygaeids, pentatomids, and coreids) (Dolling 1991).

Hemipterans are the most diverse group of hemimetabolous insects, with more than 100,000 species described (Stork 2018) and a large majority that have adopted sap-sucking life histories. Insects belonging to this guild feed on impoverished diets so that they all rely on mutualistic bacteria that live within or inside insect cells (they are termed endosymbionts) to synthesise essential nutrients that the insect cannot acquire directly from the diet (Douglas 1998). These primary, or obligate, symbionts are more like an organelle than an independent organism and are found in almost all aphids (they carry *Buchnera aphidicola*), psyllids (*Carsonella ruddii*), whiteflies (*Portiera aleyrodidarum*) and mealybugs (*Tremblaya princeps*), among others (Moran, McCutcheon & Nakabachi 2008). Such endosymbionts are required for insect survival and reproduction, transmitted from mother to offspring with high fidelity, and thought to be at the core of the diversification process of their insect hosts (Moran, McCutcheon & Nakabachi 2008). In addition to these obligate symbionts, sap-sucking insects have diverse associations with facultative (also, secondary) endosymbionts. Some of these symbionts are known to increase their hosts’ tolerance to environmental stressors like heat shocks, or to protect them against natural enemies like fungal pathogens or parasitic wasps (Oliver, Smith & Russell 2014; Guo *et al.* 2017). Although these symbionts are not required for host survival, they are nowadays recognised as fundamental to understand the ecology of sap-sucking insects, including the niche they occupy, their distribution ranges and their interactions with other members in the community. Opposite to obligate symbionts, facultative endosymbionts of sap-sucking insects are often transmitted horizontally (Caspi-Fluger *et al.* 2012; Ahmed *et al.* 2013; Chrostek *et al.* 2017) thus being mobile elements that can release herbivore populations from their natural enemies, or allow them to colonise previously inhospitable habitats. The role of facultative endosymbionts in host plant use, adaptation to abiotic conditions and interactions with natural enemies has been studied for more than a decade, particularly in well-established model systems like aphids and whiteflies (Moran, McCutcheon & Nakabachi 2008; Brownlie & Johnson 2009; Frago, Dicke & Godfray 2012; Oliver, Smith & Russell 2014; Zchori-Fein, Lahav & Freilich 2014; Zytynska & Weisser 2016; Guo *et al.* 2017; Vorburger 2018; Zytynska & Meyer 2019a).

A question that has yet to be resolved is why facultative endosymbionts are often found in only a fraction of the individuals of a population, or why they are more abundant in some populations throughout the distribution range of a given species (Zytynska & Weisser 2016). One explanation is that in the absence of the environmental stress or of natural enemy pressure, facultative symbionts can incur fitness costs (Russell & Moran 2006; Vorburger & Gouskov 2011; Vorburger, Ganesanandamoorthy & Kwiatkowski 2013). Experimental work on the potential fitness costs and benefits of hosting bacterial symbionts indicate that these effects are variable across symbiont species, host species, and also strains or genotypes within these species (Russell & Moran 2006; Rouchet & Vorburger 2012). Variation in the strength of costs and benefits across different environments can lead to the differences in infection rates we see in natural systems (Zytynska & Weisser 2016; Zytynska & Venturino 2018). Here, we present results from a meta-analysis study to understand the generality of the costs and benefits of hosting symbionts across sap-sucking insects. The majority of work in this area has been on aphids and their nine common facultative bacterial symbionts *Hamiltonella defensa*, *Regiella insecticola*, *Serratia symbiotica*, *Rickettsia*, *Ricketsiella*, *Spiroplasma*, X-type (PAXS), *Arsenophonus*, and *Wolbachia*. A number of recent reviews describe the roles these symbionts can have on their hosts, and therefore we will not provide this detail here (Oliver, Smith & Russell 2014; Zytynska & Weisser 2016; Guo *et al.* 2017; Vorburger 2018). Understanding the benefits different endosymbiont species confer to insect herbivores, and the costs they impose is thus crucial to understand one of the most diverse communities of animals in terrestrial ecosystems.

Several recent reviews have highlighted the importance of these symbionts in understanding their hosts’ biology, but currently there is a lack of a quantitative and systemic analysis of the published evidences exploring whether the different endosymbionts are actually beneficial or not. Although most research on this topic has been limited to few model systems and symbiont species, we believe the amount of data available is large enough to perform a meta-analysis. Such a systemic analysis will provide insights into the role of these microbial symbionts in different sap-sucking insect groups and will identify knowledge gaps at the level of both research questions and insect (or symbiont) taxa.

In this review we address the following questions:

1. Symbionts are widespread in sap-sucking insects, but which groups of species have been sufficiently experimentally examined for the effects of symbionts on the host?
2. Many researchers make the assumption that symbionts must incur a cost to the host (otherwise why don’t all individuals co-host multiple beneficial symbionts?); are these costs general across host and symbiont species?
3. Benefits of hosting symbionts must outweigh any costs for associations to persist, but do we have sufficient experimental data to show that symbionts confer general benefits across host and symbiont species?
4. The cost-benefit trade-off for hosting symbionts can be mediated by the immediate environment (biotic and abiotic) depending on the benefit conferred. Sap-sucking insect communities are dependent on their plant hosts, so to what extent can their host-plant species affect the costs and benefits of hosting symbionts?

## Methods

The majority of experimental work has used aphid systems, but there is growing popularity to study this in whitefly. Literature and data pertaining to other phloem-feeders was searched for, but with little to no papers relevant to our meta-analyses. We detail the individual search terms and methods below for each group.

### Whitefly meta-analysis

We searched for relevant literature using keyword searches in Web of Science finding papers published until the end of March 2018. We used the terms: whitefl* OR bemisia* OR siphoninus* OR Trialeurodes OR Aleurodicus OR Aleuronudus OR Dialeurodicus OR Metaleurodicus OR Palaealeurodicus OR Paraleyrodes AND symbio*. Resulting in 220 potential papers. To be included in the meta-analysis, papers had to satisfy the following inclusion criteria: (1) data on at least one whitefly species; (2) an experimental test of symbiont effects on traits, either experimentally cured, a comparison of field-collected infected and uninfected whitefly, or introgression of symbiont via crossing; (3) any of the following types of variables tested: any behaviour, growth, fecundity, survival, or parasitism related variable; and, (4) data on means, an estimation of variation and sample size. For the whitefly data, two studies were eliminated because the antibiotic treatment also eliminated the primary symbiont. The final dataset consisted of 9 papers (from 1993-2017, see Appendix 1 for list of included papers), and the only variables of interest with sufficient data were growth rate (development time, growth rate and longevity), fecundity, adult mass, and survival. To calculate overall symbiont effect on growth the variables development time, growth rate and longevity were pooled together. For growth rate and longevity, larger is better for the insect, but the opposite is true for development time (based on the slow-growth-high-mortality hypothesis). In the overall test for growth, yi was thus multiplied by −1.

### Aphid meta-analysis

We searched for relevant literature using keyword searches in Web of Science finding papers published until the end of 2018. We used the terms: ("aphid*" AND ("Hamiltonella" OR "Regiella" OR "Serratia" OR "Rickettsia" OR "Rickettsiella" OR "Sprioplasma" OR "Arsenophonus" OR "Wolbachia" OR "X-type" OR "PAXS")). This resulted in the extraction of 459 papers. To be included in the meta-analysis, papers had to satisfy the following inclusion criteria: (1) data on at least one aphid species (Hemiptera: Aphididae); (2) an experimental test of symbiont effects on aphids traits, either experimentally cured (Experimental) or a comparison of field-collected infected and uninfected aphids (Natural); (3) any of the following types of variables tested: aphid behaviour, growth, fecundity, survival, or parasitism-related; and, (4) data on means, an estimation of variation and sample size. This resulted in 75 potential papers, but we then included an additional inclusion criterion to include only traits with independent data points across at least three independent studies, to reduce bias from single-lab results. Further, only data where aphids hosted single symbionts were able to be used, as there was not sufficient data on multiple infections (10 studies, 37 data points [35 from *A. pisum* aphids] across 8 different symbiont combinations). In our final 57 papers (1997-2018; Figure 1a, see Appendix 2 for list of included papers) we were able to analyse the effect of bacterial symbionts on: Development (age at first reproduction), Lifespan (longevity), Mass (fresh weight), Fecundity (number of offspring), and Parasitism (proportion of aphids parasitized).

**Figure 1.**
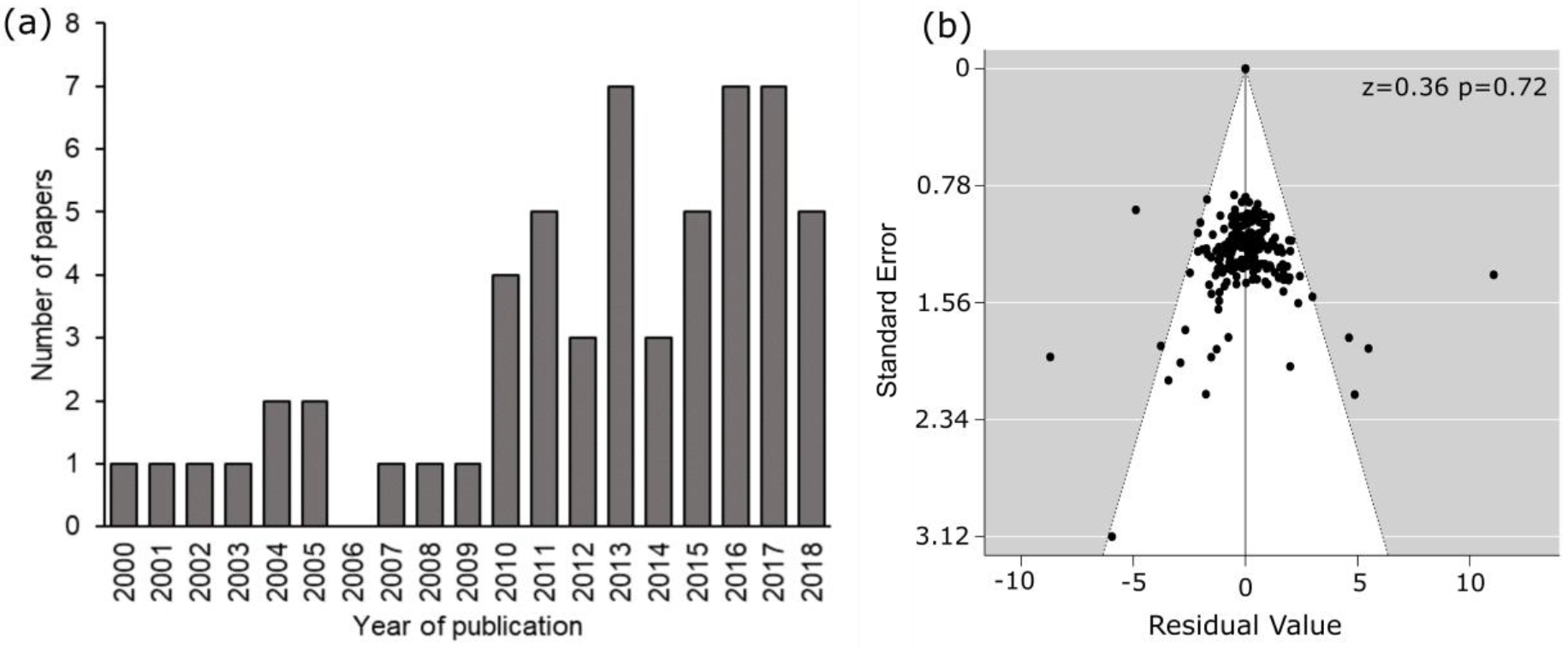
Summary of aphid meta-analysis data. (a) Number of aphid symbiont papers across year of publication for the 57 papers used in the meta-analysis, and (b) funnel plot showing no strong publication bias across these papers

To account for differences between studies that artificially cured/inoculated aphid lines, and those comparing naturally-collected infected to naturally uninfected aphid lines, we pooled data within studies across aphid genotypes and symbiont strains (i.e. removing effects of within-species genetic variation); this was necessary since no cross-comparison of common aphid or symbiont genetic lines was possible. Data was still separated within a study for aphid species, symbiont species, and host-plant (experimental plant, and plant of collection where possible).

The meta-analysis was conducted in R v3.5.1 (R Core Team 2019) in RStudio v1.1.463 (RStudio Team 2018) using the package metafor (Viechtbauer 2010). The standardised mean difference was used with unbiased estimates of the sampling variances (SMDH, giving Hedges’ g). This measure was used since it gives a direct effect size comparison of the treated (infected with a symbiont) to untreated (no symbiont control) data. We used a meta-analytic linear mixed effects model (rma) to test the effect of hosting symbionts on the different aphid traits. ‘Study’ was included as random effect to account for multiple data point across aphid and symbiont species within individual studies. Publication bias was assessed by testing the funnel-plot asymmetry (Figure 1b for overall data, Figure S1 for data subsets). Data were also subset into those where aphid lines had been directly compared through experimental curing/infecting (Experimental) or a comparison of field-collected infected and uninfected aphids (Natural), and analyses run as above on each separate dataset. The mean effect size and 95% confidence intervals are presented; the mean effect size was considered significantly different from 0 if its 95% CI did not include zero, and level of significant given from model outputs.

Further, we subset the data by aphid trait (i.e. one model for each trait) and explored differences across aphid or symbiont species within these using meta-analytic linear mixed effects models by including ‘aphid species’ and ‘symbiont species’ as moderators (equivalent to main effects in standard linear models). The interaction term was considered but in no case was there sufficient data for this to be a meaningful term to include. We used model comparisons to estimate the effect of symbiont species and aphid species using a LRT (likelihood ratio test) giving Chi-square and associated p-values. While the overall effects of different aphid and symbionts species were analysed and relevant results presented in the main text, for visual representation we present results from additional analyses that separated the aphid species data into two categories: (a) *Acyrthosiphon pisum* aphid data (the commonly-used model pea aphid species) and, (b) all other aphids (often representing less than half of the total data points). Only aphid-symbiont combinations with at least two data points are presented in the figures.

## Results

### Whitefly traits

The included studies are limited to a single species of whitefly (*B. tabaci*) and include some of its biotypes (Q, B, among others) and three symbionts: *Hamiltonella*, *Wolbachia* and *Rickettsia*. Tests are performed on different host plants and symbionts usually removed with antibiotics or via introgression (Asiimwe, Kelly & Hunter 2014). Across all symbiont species, there was a cost of carrying symbionts in *B. tabaci* through reduced fecundity (number of offspring). The following traits were not significantly affected by symbiont carrying: growth (development time (days), growth rate, longevity (days)), adult mass (body length (mm)) and survival (from egg to adult) (Figure 2a).

**Figure 2.**
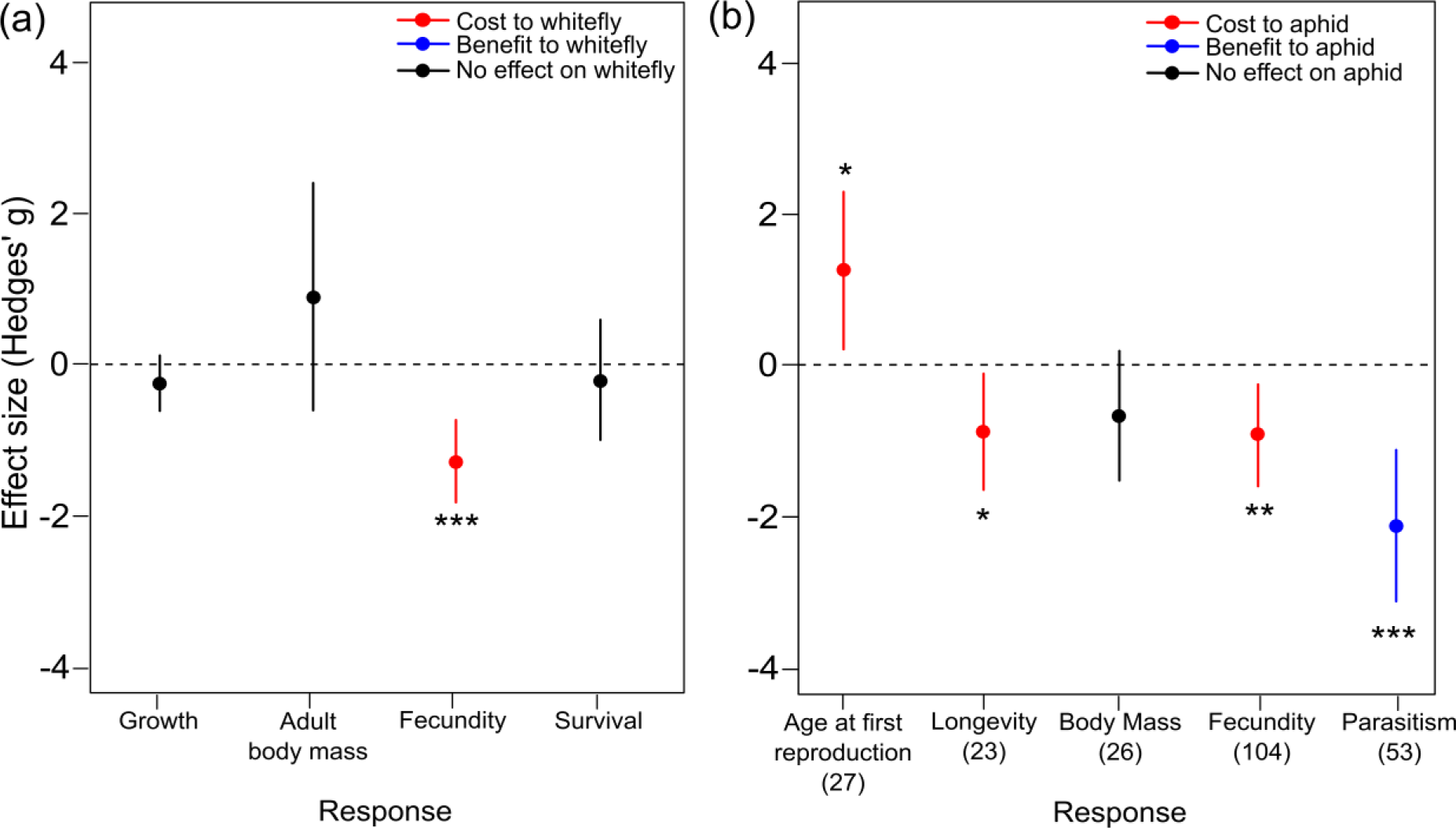
Overall effect of endosymbionts on (a) whitefly *Bemisia tabaci* across four different B. tabaci life history traits, and (b) aphids (Hemiptera: Aphididae) across five different variables, including four related to aphid life history traits and the last related to resistance to parasitism (proportion of aphids parasitized). * P<0.05, ** P<0.01, *** P<0.001

### Aphid traits

Our meta-analysis was based on data from 57 papers (2000-2018; Figure 1a, see Appendix 2 for list of included papers) yielding 233 data points from 11 aphid species: *Acyrthosiphon pisum* (n=123), *Aphis fabae* (n=32), *Sitobion avenae* (n=32), *Aphis craccivora* (n=13), *Acyrthosiphon kondoi* (n=12), *Macrosiphum euphorbiae* (n=5), *Megoura crassicauda* (n=5), *Aphis glycines* (n=4), *Myzus persicae* (n=3), *Rhopalosiphum padi* (n=3), *Obtusicauda frigidae* (n=1). We compared aphid traits of aphids infected with a symbiont to uninfected aphids (data were pooled within a study across aphid genotypes and symbiont strains, but separated by aphid species, symbiont species, and host-plant). The data were robust against publication bias as measured through funnel-plot asymmetry (z = 0.363, p = 0.717; Figure 1b).

The number of data points within our final dataset for the different response variables were: Age at first reproduction (N=27: Experimental n=26, Natural n=1; from 14 papers), Longevity (N=23: Experimental n=20, Natural n=3; from 12 papers), Mass (N=26: Experimental n=18, Natural n=8; from 15 papers), Fecundity (N=104: Experimental n=87, Natural n=17; from 40 papers), Parasitism (N=53: Experimental n=36, Natural n=17; from 28 papers).

### Effects across all aphids and symbionts

Across all aphid and symbiont species, there was a cost to aphids through increased development time (age at first reproduction), reduced longevity (days), reduced fecundity (number of offspring) (Figure 2b). However, there was also a strong benefit to aphids of hosting endosymbionts that conferred resistance against attacks by parasitic wasps (reduced proportion of aphids with a symbiont are parasitized; Figure 2b). The measures for age at first reproduction, longevity, and parasitism, were relatively consistent across studies and therefore we are able to present mean values for these effects. We found that age at first reproduction for the aphids was increased from 8.75±1.21 days (uninfected controls) to 9.06±1.21 days when hosting a symbiont, and longevity was decreased by five days when hosting a symbiont (control: 30.31±5.02 days, symbiont: 24.33±4.95 days). The proportion of aphids parasitized reduced from 0.54±0.25 (uninfected controls) to 0.36±0.29 when hosting a symbiont. The measures of body mass and fecundity varied across studies, thereby reducing our ability to provide reliable mean values for these traits.

### Experimental lines versus naturally-collected lines of aphid

When the data were compared between those aphid lines that had been experimentally infected/cured and those that were collected from the field as infected or uninfected, we see that there is substantial variation in the results (Figure S2). The unequal distribution of data points (higher number of data points for experimental lines compared to natural lines) needs to be noted here and results interpreted with this potential strong bias in mind. The results indicate that the effects are stronger and less variable within experimental studies, with a lack of overall significant results and greater range of data for ‘natural’ aphid lines collected from the field. In some cases, e.g. fecundity, the data shows a potential for a change in the direction of the result; however, further exploration of the data suggests that this may be influenced by a few individual data points owing to the lack of data across multiple species, symbionts, and laboratories.

### Effects within aphid traits by symbiont and aphid species

The age at first reproduction (days) in the aphid was increased by symbionts for all aphid species, but the magnitude varied across aphid species (Χ^2^=18.54, df=6, P<0.001), and the effect varied across the different symbiont species (Χ^2^=12.07, df=7, P=0.007). *Hamitonella defensa* symbionts did not increase aphid age at first reproduction, while *Rickettsia* sp. increased only for *A. pisum* aphids and *Serratia symbiotica* symbionts for all aphids (Figure 3a).

**Figure 3.**
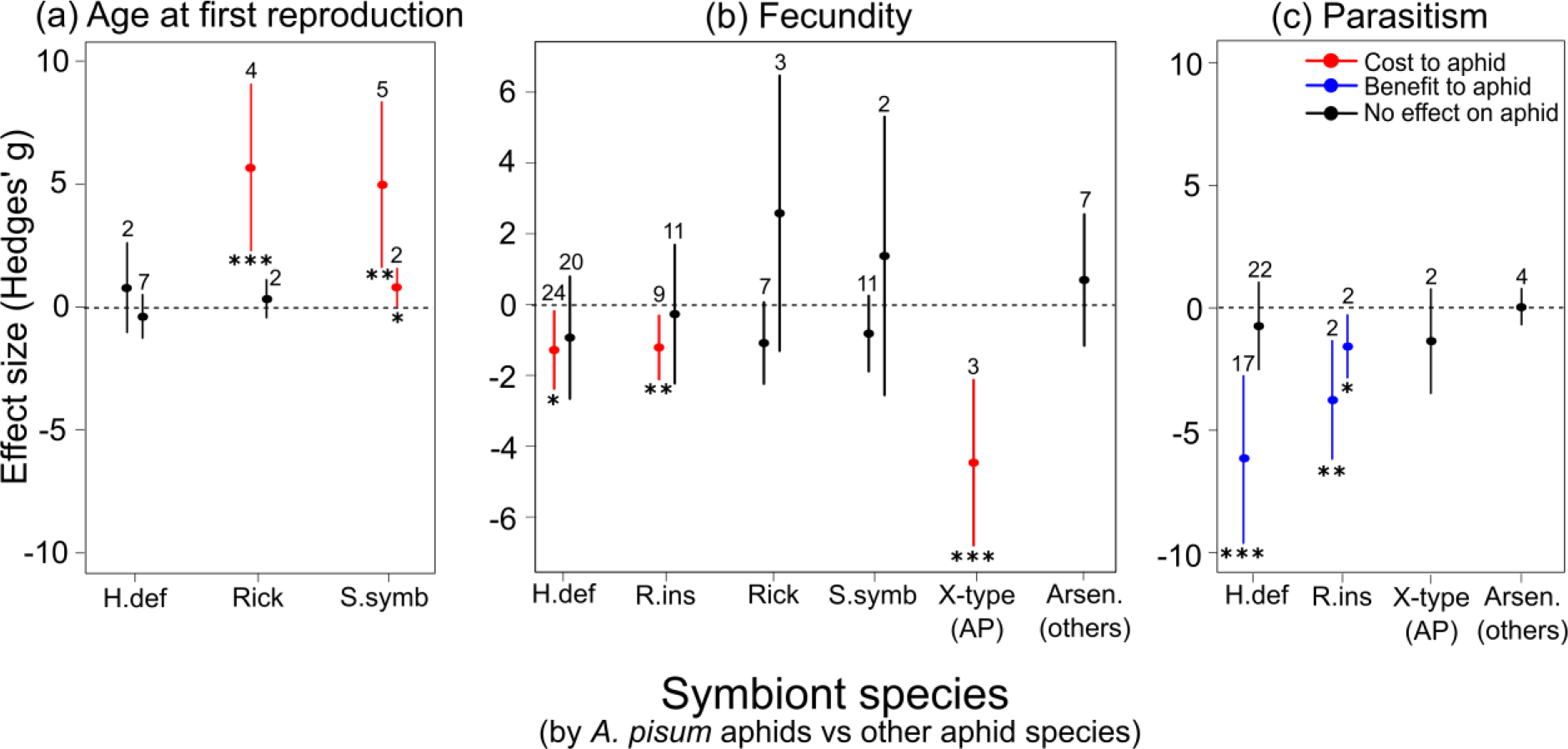
Variation in effect size of hosting symbionts for *A. pisum* pea aphids compared to other aphid species on (a) aphid age at first reproduction (days) where a positive effect indicates a lengthened development time when hosting the symbiont; (b) fecundity, where a negative effect indicates fewer offspring produced when hosting the symbiont; and, (c) parasitism, where a negative effect indicates a reduced proportion of aphids were parasitized by a parasitic wasp when hosting the symbiont. The clustered lines show results from *A. pisum* pea-aphids (left lines, AP) compared to all other aphid species combined (right lines, others). Data points are the mean effect size with 95% confidence intervals, numbers above the lines indicates the number of data points in the meta-analysis.

Overall, aphid fecundity was reduced for aphids hosting a symbiont (Figure 2b) but the effect size was dependent on the aphid species (Χ^2^=13.79, df=9, P=0.003) and, in part, the symbiont (Χ^2^=12.44, df=7, P=0.088). Both *H. defensa* and *Regiella insecticola* reduced the fecundity of *A. pisum* aphids but there was no evidence for this to affect other aphid species (Figure 3b). *Rickettsia*, *S. symbiotica*, and *Arsenophonus sp*. had no effect on aphid fecundity, while the *X-type* symbiont decreased *A. pisum* aphid fecundity (not studied in other aphids) (Figure 3b).

Symbiont-conferred resistance to parasitic wasps was variable across symbiont (Χ^2^=18.57, df=7, P=0.002) and aphid species (Χ^2^=3.73, df=7, P=0.053). *Regiella insecticola* symbionts reduced the proportion of aphids parasitized for all aphids studied, whereas *H. defensa* only reduced this for *A. pisum* aphids, no other symbionts showed a significant effect (Figure 3c).

The influence of aphid or symbiont species was not presented for the aphid traits of body mass or longevity due to strong effects of publication bias within these traits (Figure S2), potentially leading to biased conclusions; however, analysis indicated little variation in the effect of symbiont hosting across the different aphid and symbiont species tested.

## Discussion

In this meta-analysis we have explored the benefits and potential costs associated with carrying facultative endosymbionts in phloem-feeding insects. This is the first study that explores this question quantitatively across phloem-feeding Hemiptera [i.e. whiteflies, mealybugs, aphids, psyllids, planthoppers, leafhoppers, lygaeids, pentatomids and coreids (Dolling 1991)] and Thysanoptera. Our first highlight is that, although these two Orders comprise almost 110,000 described species (Stork 2018), most of them sap-suckers, information on the costs and benefits of facultative symbiont infection is limited to aphids and whiteflies. Within these two groups, we found a strong bias towards a few well-studied species, such as the model pea aphid (*A. pisum)*, the black-bean aphid (*Aphis fabae*), the cereal aphid (*Sitobion avenae*), and the whitefly *Bemisia tabaci*. This bias towards a few model species is likely due to their importance as agricultural pest species and long history of ecological studies involving these species. However, there are many other agriculturally-important sap-feeders, particularly aphid species, that have been little used in symbiont studies. Other than lack of research effort, one reason for this bias might be due to the difficulty of artificially removing symbionts in some insect species. In aphids using antibiotics to "cure" them from secondary symbionts is simple and well-documented, albeit time-consuming (Simon *et al.* 2007); however, this technique does not work for the potato aphid *Macrosiphum euphorbiae* because the antibiotic treatment eliminates both facultative and the obligatory *Buchnera* symbionts resulting in aphid death (Hackett, Karley & Bennett 2013). One work around, as done by the authors working on *M. euphorbiae*, is by testing symbiont effects in various field-collected genotypes with and without the bacterium (termed ‘Natural’ aphid lines in this meta-analysis), or creating infected lines via introgression (as done in whiteflies, e.g. (Asiimwe, Kelly & Hunter 2014)).

We find support for an overall fitness cost of hosting bacterial symbionts in the aphid and whitefly species studied; yet, the impact of these is strongly insect and symbiont species dependent. The general costs to the aphid occur through increased time until the first reproduction, reduced fecundity and reduced longevity. Thus, symbionts decrease aphid fitness by delaying development, reducing lifespan, and reducing offspring production during this time. Hosts that carry costly symbionts but that do not confer any benefit are expected to be lost in populations via purifying selection. We could show that there are general benefits of symbionts through decreased parasitism rates, such that certain symbionts protect certain aphid species from attack by specialist parasitic wasps (first shown by Oliver *et al.* 2003).

Although other benefits, such as resistance to entomopathogenic fungi or heat stress have been highlighted in reviews on aphid symbiont effects (Oliver, Smith & Russell 2014; Zytynska & Weisser 2016; Guo *et al.* 2017; Vorburger 2018), these traits lack sufficient data across multiple aphid species and symbionts to be included in a meta-analysis. In whiteflies, facultative symbionts were found to be costly only through reduced fecundity, but as far as we are aware any benefits such as symbiont-mediated resistance to natural enemies have never been tested in this insect group. Despite this, costly symbionts like *Hamiltonella* are highly prevalent in whiteflies (Gueguen *et al.* 2010; Zchori-Fein, Lahav & Freilich 2014), which suggests that the benefits associated to these bacteria are yet to be discovered.

The species-specific costs and benefits we identified in this meta-analysis have the potential to contribute to the variation in symbiont-hosting frequencies observed in the field within and among populations (Zytynska & Weisser 2016). All the experiments included in this meta-analysis compared infected with uninfected aphid lines, and in the field both infected and uninfected aphids coexist (Zytynska & Weisser 2016). The reduced fitness of aphids hosting a symbiont means they will be outcompeted by the uninfected aphids when there is no benefit (e.g. through resistance), confirming the ‘only helpful when required’ statement of Vorburger and Gouskov (2011). Yet, the magnitude of the costs and benefits will determine the impact on individual populations.

In this meta-analysis, we compared experiments that either directly assessed symbiont effects using lines of aphid that had been artificially cured or infected, or compared lines of naturally-infected to naturally-uninfected aphids collected to the field. We found that this separation resulted in strong differences in the effects. A greater amount of variation with reduced effects on the aphid were seen for those aphid lines that were collected from the field. Since this data comparison was potentially highly biased, with small sample sizes for the naturally-collected aphids, we must interpret this carefully.

However, it may suggest that ‘successful’ aphid-symbiont combinations incur fewer costs, but perhaps also reduced benefits. Or, this increased variation in effect size might indicate it is strongly dependent on the particular combination of aphid (species, genotype), symbiont (species, strain), and surrounding challenges (e.g. microclimate, host-plant, parasitism rate). Perhaps important benefits can only be appreciated under more natural conditions. A field study in the US, for example, revealed that the prevalence of the defensive symbiont *H. defensa* in *A.* pisum aphids increased throughout the season in response to increased densities of parasitic wasps (Smith *et al.* 2015). This correlation, however, was only significant in one of the two sites studied, a result that may reflect that parasitic wasps are not the only natural enemies dictating the fate of symbiont-carrying insects. In addition, most studies have been done on a restricted set of host plants, while only a small proportion of sap-sucking insects are monophagous (most feed on more than one plant species). Data were collected on ‘experimental host-plant’ and ‘host-plant of aphid collection’ but was insufficient for inclusion in the meta-analysis, indicating the need for more empirical studies in this area.

Bringing the plant layer into account will certainly change our understanding of the cost-benefit balance of symbiont infection in phloem feeders, with work suggesting the surrounding plant community can have strong impacts on aphid endosymbiont communities (Zytynska *et al.* 2016; Zytynska & Meyer 2019a). Moreover, in the aphids *A. pisum* and *C. cedri*, the facultative symbiont *S. symbiotica* can assist the obligatory symbiont at the nutritional level, potentially enabling host-feeding on a wider selection of plants (Koga, Tsuchida & Fukatsu 2003; Lamelas *et al.* 2011). In whiteflies (Su *et al.* 2015) and aphids (Frago *et al.* 2017) recent studies also revealed that symbionts are able to help their hosts circumvent plant defences that are triggered upon insect attack. Altogether, this means that symbionts could have a much wider impact on aphid populations than is currently empirically tested. Thus, to better understand the balance between costs and benefits of symbiont infection, and ultimately their prevalence in natural populations, a wider community perspective is necessary. A recent review discusses the importance of the immediate surrounding biotic community (plant diversity, natural enemy diversity) in combination with the abiotic environment on mediating aphid-symbiont interactions (Zytynska & Meyer 2019a).

Complex community interactions also occur among symbionts inside their hosts, with potential implications for the benefits that these bacteria provide (Ferrari & Vavre 2011). Our meta-analysis is based only on insects with single symbiont infections since we lacked sufficient data on the role of multiple symbiont infections. In the field it is estimated that aphids, for instance, host 0-4 symbionts per individual (Ferrari *et al.* 2012; Russell *et al.* 2013; Smith *et al.* 2015; Zytynska *et al.* 2016), and that multiple infections are particularly common in some genus like *Macrosiphum* (Henry et al. 2015). More importantly, where symbiont co-infections occur strong fitness costs are often observed (Oliver, Moran & Hunter 2006; Guay *et al.* 2009; Leclair *et al.* 2017; McLean *et al.* 2018). It would be very interesting to test this across different aphid species, asking whether the proportion of multiple infections correlates negatively with the fitness costs they impose. To better understand how multiple infections arise, more work on how facultative symbionts are horizontally transferred within populations of the same species or among species is also needed. While aphid symbionts are predominantly vertically transmitted from mother to offspring, there is also evidence of horizontal transfer of symbionts among aphids during sexual reproduction (Moran & Dunbar 2006), by parasitoids when ovipositing eggs into aphids (Gehrer & Vorburger 2012), or even through infected honeydew (Darby & Douglas 2003). Based on a few laboratory studies, aphid hosts impose little constrains to symbiont acquisition even if the host already carries a facultative symbiont, demonstrated by successful microinjecting of different symbiont species and strains into pea aphids (Leclair *et al.* 2017; McLean *et al.* 2018). A recent paper used mathematical modelling to further show the importance of horizontal transmission of symbionts among aphids, focusing on the potential of parasitoid wasps to transmit protective symbionts among aphids (Zytynska & Venturino 2018). A low rate of horizontal transmission led to coexistence of uninfected aphids, infected aphids, and parasitoid wasps, with the percentage of infected aphids ranging from 30-70% which is in agreement with infection rates observed in field surveys (Zytynska & Weisser 2016; Zytynska & Venturino 2018).

Many of these symbionts protect agricultural insect pests from their natural enemies, which is counteractive to the aims of biological control programs (Vorburger 2018). Understanding how different insect species interact with the common symbionts species, how symbionts are transmitted between insect individuals, and the likely costs and benefits, all help to devise management programs to reduce the impact of these symbionts. In a closed greenhouse system, the spread of a protective symbiont can hinder biological control efforts, yet strategies to increase natural enemy diversity could mitigate this impact (Vorburger 2018). In the field, if natural enemy density is low, as might occur in a monoculture crop field, then insects with no symbionts (higher fitness) would outcompete insects with symbionts (Zytynska & Meyer 2019b). Coupled with pesticide resistance evolution (e.g. seen in cereal aphids in Europe; Malloch *et al.* (2016)), this can lead to increased risks of pest outbreaks.

## Conclusion

Current molecular methods allow us to study the intricate ways insects establish mutualistic symbioses with microbial partners. We used meta-analysis techniques to show the general costs (through increased development time, reduced longevity and reduced fecundity) and benefits (increases resistance to parasitic wasps) of hosting bacterial symbionts in sap-sucking insects. Current data is strongly biased towards a few species of aphid and whitefly, and that there is a large variation of effects among insect as well as symbiont species. Thus, the results cannot reliably be extrapolated to other phloem-feeding taxa, and not even to other aphid or whitefly species. The impact of cost-benefit trade-offs in natural systems are still to be uncovered, but an appreciation of the diversity of potential outcomes due to the species or genetics of the insect/symbiont and the surrounding environment (plant diversity, natural enemy diversity, microclimate) will benefit the design of future studies. While many of the studied insect species are agricultural pests, studies in which the phenotypic consequences of facultative symbiont infection are measured in non-model species are urgently needed. For agricultural systems, the spread of protective symbionts in sap-sucking insects can hinder biological control efforts while reduced densities of natural enemies might select for uninfected aphids with higher reproductive fitness. In both cases, this can lead to pest outbreaks.

## Data availability

All papers used in the meta-analysis are detailed in the appendices. Data will be made readily available through contact with the corresponding author until final publication of the paper when data will be made publically available.

## Acknowledgements

This project is the result of one of one of the work packages of the EU COST action FA1405 on Crop-arthropod-microbe interactions (www.cost-camo.eu). Special thanks to Arjen Biere and Alison Bennett. SEZ was supported by a British Ecological Society small research grant (SR16/1069). EF was funded by the Regional Council of Reunion, the Departmental Council of the Region Reunion, the European Union (EAFRD) and CIRAD.

