## Supplementary Material for "A meta-analysis on the benefits and costs of hosting secondary endosymbionts in sap-sucking insects"

**Figure S1. Funnel plots and test statistics for potential publication bias across subsets of the data**

**Figure S2. Effect of hosting a symbiont on aphid traits, separated by study method.**

**Appendix 1. List of included studies in the whitefly analysis**

**Appendix 2. List of included studies in the aphid analysis**

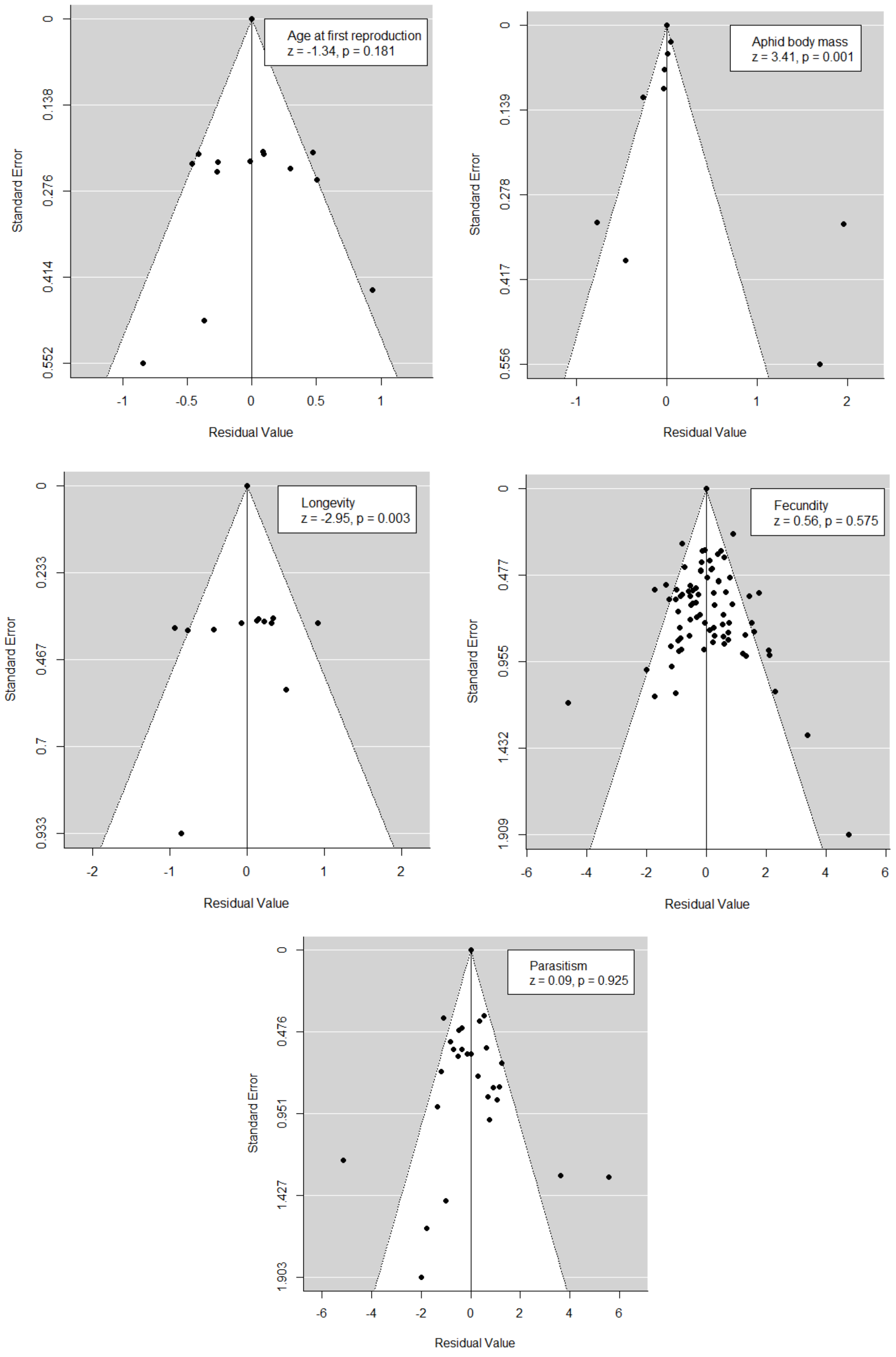


**Figure S2. Funnel plots and test statistics for potential publication bias across subsets of the data** (by aphid traits). P<0.05 indicates potential publication bias. Only ‘Age at first reproduction’, ‘Fecundity’, and ‘Parasitism’ are explored further within the subset of data as they show no publication bias.

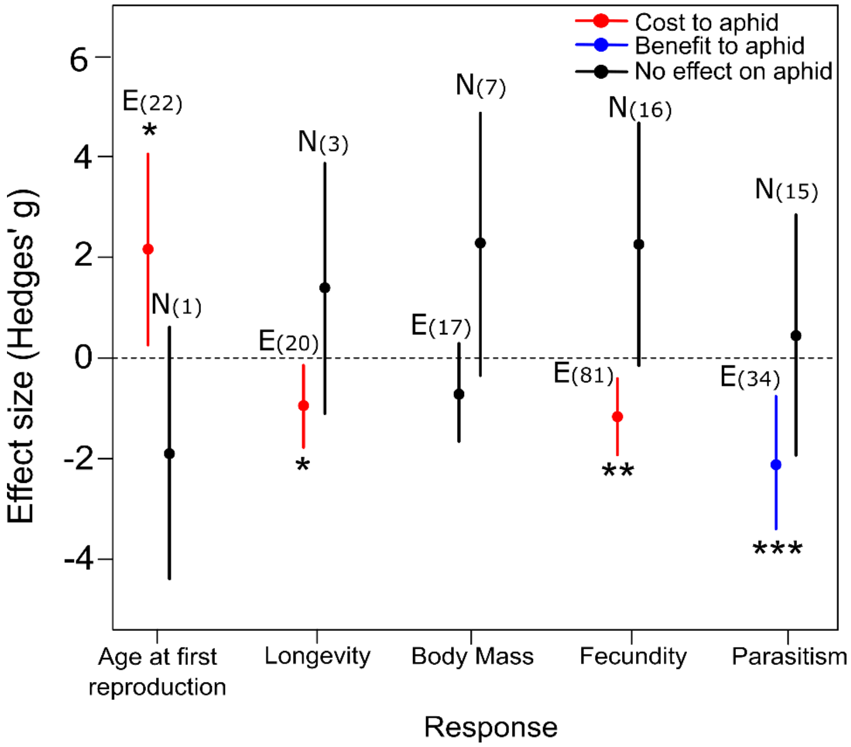


**Figure S2. Effect of hosting a symbiont on aphid traits, separated by study method.** Experimental aphids (E, left line for each aphid trait) are lines where aphids were artificially cured or infected and lines directly compared, whereas naturally-collected aphids (N, right line for each aphid trait) compared aphids collected from the field that were naturally uninfected to those that were naturally infected by symbionts. The numbers in brackets next to the N or E indicate the number of data points included in the analysis.

* P<0.05, ** P<0.01, *** P<0.001

**Appendix 1. List of included studies in the whitefly analysis (1-8)**

**Appendix 2. List of included studies in the aphid analysis (1-57)**


1. Alkhedir H, Karlovsky P, Mashaly AMA, & Vidal S (2016) Specialization and host plant use of the common clones of Sitobion avenae (Homoptera: Aphididae). *Applied Entomology and Zoology* 51(2):289-295.
2. Barribeau SM, Sok D, & Gerardo NM (2010) Aphid reproductive investment in response to mortality risks. *Bmc Evolutionary Biology* 10.
3. Castaneda LE, Sandrock C, & Vorburger C (2010) Variation and covariation of life history traits in aphids are related to infection with the facultative bacterial endosymbiont Hamiltonella defensa. *Biological Journal of the Linnean Society* 100(1):237-247.
4. Cayetano L, Rothacher L, Simon JC, & Vorburger C (2015) Cheaper is not always worse: strongly protective isolates of a defensive symbiont are less costly to the aphid host. *Proceedings of the Royal Society B-Biological Sciences* 282(1799).
5. Cayetano L & Vorburger C (2013) Effects of Heat Shock on Resistance to Parasitoids and on Life History Traits in an Aphid/Endosymbiont System. *Plos One* 8(10).
6. Cayetano L & Vorburger C (2013) Genotype-by-genotype specificity remains robust to average temperature variation in an aphid/endosymbiont/parasitoid system. *Journal of Evolutionary Biology* 26(7):1603-1610.
7. Cayetano L & Vorburger C (2015) Symbiont-conferred protection against Hymenopteran parasitoids in aphids: how general is it? *Ecological Entomology* 40(1):85-93.
8. Chen DQ, Montllor CB, & Purcell AH (2000) Fitness effects of two facultative endosymbiotic bacteria on the pea aphid, Acyrthosiphon pisum, and the blue alfalfa aphid, A-kondoi. *Entomologia Experimentalis Et Applicata* 95(3):315-323.
9. Clarke HV, Cullen D, Hubbard SF, & Karley AJ (2017) Susceptibility of Macrosiphum euphorbiae to the parasitoid Aphidius ervi: larval development depends on host aphid genotype. *Entomologia Experimentalis Et Applicata* 162(2):148-158.
10. Clarke HV, Foster SP, Oliphant L, Waters EW, & Karley AJ (2018) Co-occurrence of defensive traits in the potato aphid Macrosiphum euphorbiae. *Ecological Entomology* 43(4):538-542.
11. Dion E, Polin SE, Simon JC, & Outreman Y (2011) Symbiont infection affects aphid defensive behaviours. *Biology Letters* 7(5):743-746.
12. Doremus MR & Oliver KM (2017) Aphid Heritable Symbiont Exploits Defensive Mutualism. *Applied and Environmental Microbiology* 83(8).
13. Dykstra HR*, et al.* (2014) Factors Limiting the Spread of the Protective Symbiont Hamiltonella defensa in Aphis craccivora Aphids. *Applied and Environmental Microbiology* 80(18):5818-5827.
14. Erickson DM, Wood EA, Oliver KM, Billick I, & Abbot P (2012) The Effect of Ants on the Population Dynamics of a Protective Symbiont of Aphids, Hamiltonella defensa. *Annals of the Entomological Society of America* 105(3):447-453.
15. Ferrari J, Scarborough CL, & Godfray HCJ (2007) Genetic variation in the effect of a facultative symbiont on host-plant use by pea aphids. *Oecologia* 153(2):323-329.
16. Fukatsu T, Tsuchida T, Nikoh N, & Koga R (2001) Spiroplasma symbiont of the pea aphid, Acyrthosiphon pisum (Insecta : Homoptera). *Applied and Environmental Microbiology* 67(3):1284-1291.
17. Hansen AK, Vorburger C, & Moran NA (2012) Genomic basis of endosymbiont-conferred protection against an insect parasitoid. *Genome Research* 22(1):106-114.
18. Heyworth ER & Ferrari J (2015) A facultative endosymbiont in aphids can provide diverse ecological benefits. *Journal of Evolutionary Biology* 28(10):1753-1760.
19. Heyworth ER & Ferrari J (2016) Heat Stress Affects Facultative Symbiont-Mediated Protection from a Parasitoid Wasp. *Plos One* 11(11).
20. Katayama N, Tsuchida T, Hojo MK, & Ohgushi T (2013) Aphid Genotype Determines Intensity of Ant Attendance: Do Endosymbionts and Honeydew Composition Matter? *Annals of the Entomological Society of America* 106(6):761-770.
21. Koga R, Tsuchida T, & Fukatsu T (2003) Changing partners in an obligate symbiosis: a facultative endosymbiont can compensate for loss of the essential endosymbiont Buchnera in an aphid. *Proceedings of the Royal Society B-Biological Sciences* 270(1533):2543-2550.
22. Leclair M*, et al.* (2017) Consequences of coinfection with protective symbionts on the host phenotype and symbiont titres in the pea aphid system. *Insect Science* 24(5):798-808.
23. Leclair M*, et al.* (2016) Diversity in symbiont consortia in the pea aphid complex is associated with large phenotypic variation in the insect host. *Evolutionary Ecology* 30(5):925-941.
24. Lenhart PA & White JA (2017) A defensive endosymbiont fails to protect aphids against the parasitoid community present in the field. *Ecological Entomology* 42(5):680-684.
25. Leonardo TE (2004) Removal of a specialization-associated symbiont does not affect aphid fitness. *Ecology Letters* 7(6):461-468.
26. Leybourne DJ, Bos JI, Valentine TA, & Karley AJJIS (2018) The price of protection: a defensive endosymbiont impairs nymph growth in the bird cherry-oat aphid, Rhopalosiphum padi.
27. Li SR*, et al.* (2018) Effects of a presumably protective endosymbiont on life-history characters and their plasticity for its host aphid on three plants. *Ecology and Evolution* 8(24):13004-13013.
28. Lukasik P, Dawid MA, Ferrari J, & Godfray HCJ (2013) The diversity and fitness effects of infection with facultative endosymbionts in the grain aphid, Sitobion avenae. *Oecologia* 173(3):985-996.
29. Lukasik P, Guo H, Van Asch M, Ferrari J, & Godfray HCJ (2013) Protection against a fungal pathogen conferred by the aphid facultative endosymbionts Rickettsia and Spiroplasma is expressed in multiple host genotypes and

species and is not influenced by co-infection with another symbiont. *Journal of Evolutionary Biology* 26(12):2654-2661.
30. Lukasik P, Hancock EL, Ferrari J, & Godfray HCJ (2011) Grain aphid clones vary in frost resistance, but this trait is not influenced by facultative endosymbionts. *Ecological Entomology* 36(6):790-793.
31. Luo C*, et al.* (2017) Ecological impact of a secondary bacterial symbiont on the clones of Sitobion avenae (Fabricius) (Hemiptera: Aphididae). *Scientific Reports* 7.
32. Martinez AJ, Kim KL, Harmon JP, & Oliver KM (2016) Specificity of Multi-Modal Aphid Defenses against Two Rival Parasitoids. *Plos One* 11(5).
33. Martinez AJ, Weldon SR, & Oliver KM (2014) Effects of parasitism on aphid nutritional and protective symbioses. *Molecular Ecology* 23(6):1594-1607.
34. McLean AHC & Godfray HCJ (2015) Evidence for specificity in symbiont-conferred protection against parasitoids. *Proceedings of the Royal Society B-Biological Sciences* 282(1811).
35. McLean AHC & Godfray HCJ (2017) The outcome of competition between two parasitoid species is influenced by a facultative symbiont of their aphid host. *Functional Ecology* 31(4):927-933.
36. McLean AHC, van Asch M, Ferrari J, & Godfray HCJ (2011) Effects of bacterial secondary symbionts on host plant use in pea aphids. *Proceedings of the Royal Society B-Biological Sciences* 278(1706):760-766.
37. Montllor CB, Maxmen A, & Purcell AH (2002) Facultative bacterial endosymbionts benefit pea aphids Acyrthosiphon pisum under heat stress. *Ecological Entomology* 27(2):189-195.
38. Niepoth N, Ellers J, & Henry LM (2018) Symbiont interactions with non-native hosts limit the formation of new symbioses. *Bmc Evolutionary Biology* 18:12.
39. Nyabuga FN, Outreman Y, Simon JC, Heckel DG, & Weisser WW (2010) Effects of pea aphid secondary endosymbionts on aphid resistance and development of the aphid parasitoid Aphidius ervi: a correlative study. *Entomologia Experimentalis Et Applicata* 136(3):243-253.
40. Oliver KM, Campos J, Moran NA, & Hunter MS (2008) Population dynamics of defensive symbionts in aphids. *Proceedings of the Royal Society B-Biological Sciences* 275(1632):293-299.
41. Rothacher L, Ferrer-Suay M, & Vorburger C (2016) Bacterial endosymbionts protect aphids in the field and alter parasitoid community composition. *Ecology* 97(7):1712-1723.
42. Rouchet R & Vorburger C (2012) Strong specificity in the interaction between parasitoids and symbiont-protected hosts. *Journal of Evolutionary Biology* 25(11):2369-2375.
43. Russell JA & Moran NA (2005) Horizontal transfer of bacterial symbionts: Heritability and fitness effects in a novel aphid host. *Applied and Environmental Microbiology* 71(12):7987-7994.
44. Sakurai M, Koga R, Tsuchida T, Meng XY, & Fukatsu T (2005) Rickettsia symbiont in the pea aphid Acyrthosiphon pisum: Novel cellular tropism, effect on host fitness, and interaction with the essential symbiont Buchnera. *Applied and Environmental Microbiology* 71(7):4069-4075.
45. Skaljac M, Kirfel P, Grotmann J, & Vilcinskas A (2018) Fitness costs of infection with Serratia symbiotica are associated with greater susceptibility to insecticides in the pea aphid Acyrthosiphon pisum. *Pest Management Science* 74(8):1829-1836.
46. Tsuchida T, Koga R, Fujiwara A, & Fukatsu T (2014) Phenotypic Effect of "Candidatus Rickettsiella viridis," a Facultative Symbiont of the Pea Aphid (Acyrthosiphon pisum), and Its Interaction with a Coexisting Symbiont. *Applied and Environmental Microbiology* 80(2):525-533.
47. Tsuchida T, Koga R, & Fukatsu T (2004) Host plant specialization governed by facultative symbiont. *Science* 303(5666):1989-1989.
48. Tsuchida T, Koga R, Matsumoto S, & Fukatsu T (2011) Interspecific symbiont transfection confers a novel ecological trait to the recipient insect. *Biology Letters* 7(2):245-248.
49. Vorburger C, Ganesanandamoorthy P, & Kwiatkowski M (2013) Comparing constitutive and induced costs of symbiont-conferred resistance to parasitoids in aphids. *Ecology and Evolution* 3(3):706-713.
50. Vorburger C, Gehrer L, & Rodriguez P (2010) A strain of the bacterial symbiont Regiella insecticola protects aphids against parasitoids. *Biology Letters* 6(1):109-111.
51. Vorburger C & Gouskov A (2011) Only helpful when required: a longevity cost of harbouring defensive symbionts. *Journal of Evolutionary Biology* 24(7):1611-1617.
52. Vorburger C & Rouchet R (2016) Are aphid parasitoids locally adapted to the prevalence of defensive symbionts in their hosts? *Bmc Evolutionary Biology* 16.
53. Vorburger C, Sandrock C, Gouskov A, Castaneda LE, & Ferrari J (2009) Genotypic variation and the role of defensive endosymbionts in an all-parthogenetic host-parasitoid interaction. *Evolution* 63(6):1439-1450.
54. Wagner SM*, et al.* (2015) Facultative endosymbionts mediate dietary breadth in a polyphagous herbivore. *Functional Ecology* 29(11):1402-1410.
55. Wang D*, et al.* (2016) Comparison of fitness traits and their plasticity on multiple plants for Sitobion avenae infected and cured of a secondary endosymbiont. *Scientific Reports* 6.
56. Wulff JA, Buckman KA, Wu KM, Heimpel GE, & White JA (2013) The Endosymbiont Arsenophonus Is Widespread in Soybean Aphid, Aphis glycines, but Does Not Provide Protection from Parasitoids or a Fungal Pathogen. *Plos One* 8(4).
57. Zepeda-Paulo F, Villegas C, & Lavandero B (2017) Host genotype-endosymbiont associations and their relationship with aphid parasitism at the field level. *Ecological Entomology* 42(1):86-95.
